# Isoforms of the transcriptional cofactor SIN3 differentially regulate genes necessary for energy metabolism and cell survival

**DOI:** 10.1101/2021.12.31.474661

**Authors:** Anindita Mitra, Linh Vo, Imad Soukar, Ashlesha Chaubal, Miriam L. Greenberg, Lori A. Pile

**Affiliations:** Department of Biological Sciences, Wayne State University, Detroit, Michigan 48202; Integrative Program for Biological and Genome Sciences, University of North Carolina, Chapel Hill, NC 27599

**Author notes:** To whom correspondence should be addressed: Lori A. Pile: Department of Biological Sciences, Wayne State University, Detroit, MI 48202.

**Keywords:** SIN3, transcription, energy metabolism, mitochondrial membrane potential, mitochondrial dysfunction, electron transport chain, oxidative stress, apoptosis

## Abstract

The SIN3 scaffolding protein is a conserved transcriptional regulator known to fine-tune gene expression. In Drosophila, there are two major isoforms of SIN3, SIN3 220 and SIN3 187, which each assemble into multi-subunit histone modifying complexes. The isoforms have distinct developmental expression patterns and non-redundant functions. Gene regulatory network analyses indicate that both isoforms affect genes encoding proteins in pathways such as the cell cycle and cell morphogenesis. Interestingly, the SIN3 187 isoform uniquely regulates a subset of pathways including post-embryonic development, phosphate metabolism and apoptosis. Target genes in the phosphate metabolism pathway include nuclear-encoded mitochondrial genes coding for proteins responsible for oxidative phosphorylation. Here, we investigate the physiological effects of SIN3 isoforms on energy metabolism and cell survival. We find that ectopic expression of SIN3 187 represses expression of several nuclear-encoded mitochondrial genes affecting production of ATP and generation of reactive oxygen species (ROS). Forced expression of SIN3 187 also activates several pro-apoptotic and represses a few anti-apoptotic genes. In the SIN3 187 expressing cells, these gene expression patterns are accompanied with an increased sensitivity to paraquat-mediated oxidative stress. These findings indicate that SIN3 187 influences the regulation of mitochondrial function, apoptosis and oxidative stress response in ways that are dissimilar from SIN3 220. The data suggest that the distinct SIN3 histone modifying complexes are deployed in different cellular contexts to maintain homeostasis.

## Introduction

Cells are frequently exposed to internal and external stressors that affect their survival. Several processes allow cells to respond to such stresses and return to homeostatic conditions. Epigenetic regulators comprise an important class of proteins that contribute to maintenance of homeostasis by regulating gene expression (1). Histone lysine acetyltransferases and histone deacetylases (HDACs) function as multi-protein complexes that regulate gene expression by modifying the level of acetylation of histone tails (2). The SIN3 complex is one such complex. The SIN3 complex consists of the scaffolding protein SIN3 and its protein interacting partners. This includes histone deacetylase 1 (HDAC1) as well as distinct sets of accessory proteins, dependent on cell type (3). The SIN3 complex regulates and fine-tunes the expression of target genes rather than switching its targets on or off (4). While a few organisms, such as budding yeast, have a single SIN3 isoform, others express more than one isoform, including SIN3A and SIN3B, encoded by separate genes, in mammalian cells (5). In *Drosophila melanogaster*, two dominant isoforms of SIN3 are expressed, SIN3 220 and SIN3 187 (6–7). These isoforms are encoded by the single *Sin3A* gene and are splice variants. Across species, the SIN3 complex influences expression of genes encoding proteins that regulate multiple essential cellular pathways including those in the cell cycle and energy production (3).

Included in the gene targets of SIN3 are those encoding proteins needed for proper mitochondrial function. In yeast, *sin3* null mutants demonstrate a slow growth phenotype when grown on fermentable media and are unable to grow on non-fermentable media (8). Additionally, yeast cells lacking *SIN3* have low ATP production during log and stationary growth phases. In Drosophila S2 cultured cells, reduction of SIN3 220 results in an increase in both mitochondrial mass and number, relative to control samples (9). When SIN3 220 deficient cells are placed in low nutrient media, where they are forced to rely mainly on oxidative phosphorylation, they are unable to produce levels of ATP similar to those in control cells (8). Deletion of SIN3A in murine embryonic fibroblasts (MEFs) results in altered expression of various nuclear-encoded mitochondrial genes involved in mitochondrial respiration and metabolism (10). SIN3A deficiency in MEFs inhibits progression through the cell cycle, affects DNA replication, and causes increased apoptosis.

SIN3 and its associated HDAC regulate the response to oxidative stress in diverse organisms. Loss of Sin3 in worms results in increased reactive oxygen species (ROS) due to the misregulation of genes required to neutralize ROS levels (11). In Drosophila cultured cells, both SIN3 220 and dKDM5/LID, a histone demethylase that associates with SIN3 220, regulate expression of oxidative stress response genes (12). When Drosophila cells deficient for SIN3 220 are exposed to an external oxidative stressor, several cell cycle genes are misregulated. These changes in gene expression are coupled with an inhibition of cell cycle progression. The dependence on SIN3 to facilitate a response to oxidative stress was also observed in adult flies (13). The loss of the dominant isoform in adult flies, SIN3 187, leads to increased sensitivity to the ROS inducer paraquat along with alterations in expression of glutathione metabolism genes. In cancer cells, HDACs influence response to stress and apoptosis by regulating expression of associated genes. For instance, in human lung cancer cells, two pro-apoptotic genes, *p53* and *BAX*, are repressed by overexpression of HDACs 1, 2 and 3 (14). HDACs also regulate the acetylation of p53, thereby controlling activity and stability of the protein (15). In mice, the loss of *Sin3A* in cells of the inner cell mass results in an increased incidence of apoptosis via a p53-independent mechanism (16). Together, these results underscore the importance of SIN3 and HDACs in response to oxidative stress and apoptotic cell death.

The Drosophila SIN3 isoforms exhibit a differential pattern of expression in different tissues and during embryogenesis (17). They each associate with HDAC1, but the accessory proteins vary dependent on which SIN3 isoform is acting as the scaffolding factor (18). Both isoforms share interacting partners ARID4B, SAP130, ING1 and SDS3, while the SIN3 220 complex also contains a few unique partners such as the demethylase dKDM5/LID, EMSY and Caf1-p55. Previous studies in Drosophila have shown that loss of the dominant isoform at different stages of development results in loss of viability (13, 17). Intriguingly, while SIN3 220 can rescue loss of viability in *Sin3A* mutant flies, SIN3 187 lacks this ability (18). Although both Drosophila SIN3 isoforms regulate genes of pathways such as metabolism and oxidative stress (12, 19), relative to the role of SIN3 220 in regulating these processes, little is known about processes specifically linked to regulation by SIN3 187.

In this study, we aimed to investigate the physiological influence of different SIN3 isoforms on energy metabolism and survival through control of mitochondrial function and apoptosis. To do this, we used Drosophila cultured cells wherein the cells express SIN3 220 or were induced to ectopically express SIN3 187. Our results indicate that SIN3 187 uniquely repressed multiple nuclear-encoded mitochondrial genes that code for several electron transport chain subunits. The gene expression changes were accompanied by altered mitochondrial function, including increased proton leak and reduced coupling efficiency in cells expressing SIN3 187 compared to those with SIN3 220. Further, cells overexpressing SIN3 187 were highly sensitive to external oxidative stress, which can be attributed to the upregulation of multiple pro-apoptotic genes combined with sub-optimal mitochondrial function. These gene expression and phenotypic changes were absent in cells that express SIN3 220. Taken together, our findings suggest that distinct SIN3 isoform complexes can facilitate different physiologically relevant cellular responses that stem from unique gene expression regulatory networks.

## Results

### SIN3 187 represses several nuclear-encoded mitochondrial genes

Our laboratory was the first to investigate the differential binding of the Drosophila SIN3 isoforms and consequently, to identify pathways regulated by the isoforms. To achieve this, Drosophila S2 cells were used to perform ChIP-seq and RNA-seq followed by integration of both genome wide data sets to identify common and unique pathways controlled by SIN3 220 and SIN3 187 (19). For the ChIP-seq analysis, cells ectopically expressing either epitope tagged SIN3 220 or SIN3 187 were used to identify binding sites. To identify the genes regulated by SIN3 220, RNA interference (RNAi) was used to knockdown *Sin3A* in Drosophila S2 cells, which predominantly express SIN3 220 (19, 20). Genes that changed in expression with the knockdown were interpreted as those regulated by SIN3 220. Interestingly, in cultured cells, the ectopic expression of SIN3 187 results in a strong reduction in levels of endogenous SIN3 220, along with the replacement of SIN3 220 by SIN3 187 at many genomic loci (19). Specifically, ectopic expression of SIN3 187 leads to reduction in the levels of SIN3 220 at the RNA and protein levels (21). Cells ectopically expressing SIN3 187 were thus used to find genes regulated and bound by SIN3 187 as compared to wild type S2 cells. Some pathways regulated by both isoforms include cell cycle, metabolism and cell morphogenesis. The SIN3 187 complex also regulates unique pathways such as phosphate metabolism, apoptosis and endocytosis, demonstrating differential gene regulation by the distinct SIN3 isoforms. Specifically, published RNA-seq data showed that a number of nuclear-encoded mitochondrial genes did not change in expression following the loss of SIN3 220 while their levels were altered upon ectopic expression of SIN3 187 (12, 19). To dissect the roles of the distinct SIN3 isoforms in regulating mitochondrial function, we initially focused on this set of differentially regulated target genes (Table I) (12, 19). All genes listed, except for *MgstI*, are exclusively regulated by SIN3 187. All genes, barring two, are repressed by this SIN3 isoform.

**Table I:** Nuclear-encoded mitochondrial genes are differentially regulated by SIN3 isoforms. Genes that are both bound by SIN3 and altered in expression following perturbation in the level of the specific isoform are considered direct targets. The direction of regulation (activated or repressed) is indicated. Indirectly regulated genes are those that change in expression with the perturbation of the SIN3 isoform but the isoform was not found to bind at the promoter. Data obtained by analyzing published data sets (12, 19).

| Functional category | Gene name | Regulated by SIN3 220 | Regulated by SIN3 187 |
| --- | --- | --- | --- |
| Complex I | *ND-13B* | No | Repressed |
|  | *ND-B17* | No | Repressed |
|  | *ND-23* | No | Repressed |
|  | *ND-51* | No | Repressed |
|  | *ND-B14.5B* | No | Repressed |
|  | *ND-B16.6* | No | Indirect repression |
|  | *ND-B17.2* | No | Indirect repression |
|  | *ND-49* | No | Indirect repression |
| Complex IV | *Cype* | No | Repressed |
|  | *Scox* | No | Repressed |
|  | *CoVb* | No | Indirect repression |
| Complex V | *ATPsyn-gamma* | No | Activated |
|  | *Atpalpha* | No | Repressed |
|  | *ATPsynbeta* | No | Repressed |
|  | *ATPsynD* | No | Repressed |
|  | *ATPsynC* | No | Repressed |
|  | *ATPsynO/Oscp* | No | Repressed |
| Miscellaneous | *Cyt-c-p* | No | Repressed |
|  | *Sod2* | No | Repressed |
|  | *TFAM* | No | Repressed |
|  | *Mgstl* | Indirect activation | Activated |

To validate the differential gene regulation by the SIN3 isoforms, we utilized our previously established cultured Drosophila S2 cells, with predominant expression of either SIN3 220 or SIN3 187. To study the effect of SIN3 220 on the expression of target genes, SIN3 220 levels were reduced by knockdown of *Sin3A* using RNAi (Fig. 1*A*). To study the role of SIN3 187, we used cells carrying a stable integration of a transgene encoding SIN3 187 tagged with an HA epitope (SIN3 187HA cells). This transgene is regulated by a copper sulfate (CuSO_4_) inducible metallothionine promoter. In the absence of CuSO_4_, we detected approximately equal levels of endogenous SIN3 220 and of SIN3 187HA, due to leaky expression from the inducible promoter (Fig. 1*A*). As noted in our earlier publications, the induced expression of SIN3 187 results in nearly undetectable levels of the SIN3 220 isoform (Supp fig 1) (19, 21). We note that the addition of CuSO_4_ results in a high level of expression of SIN3 187HA, more than the level of endogenous SIN3 220 in S2 cells. While the level of expression of SIN3 187HA is higher than that of SIN3 220, the number of promoters bound and the level of binding at those regulated promoters is similar comparing the two isoforms (Supp fig 2) (19).

**Figure 1.**
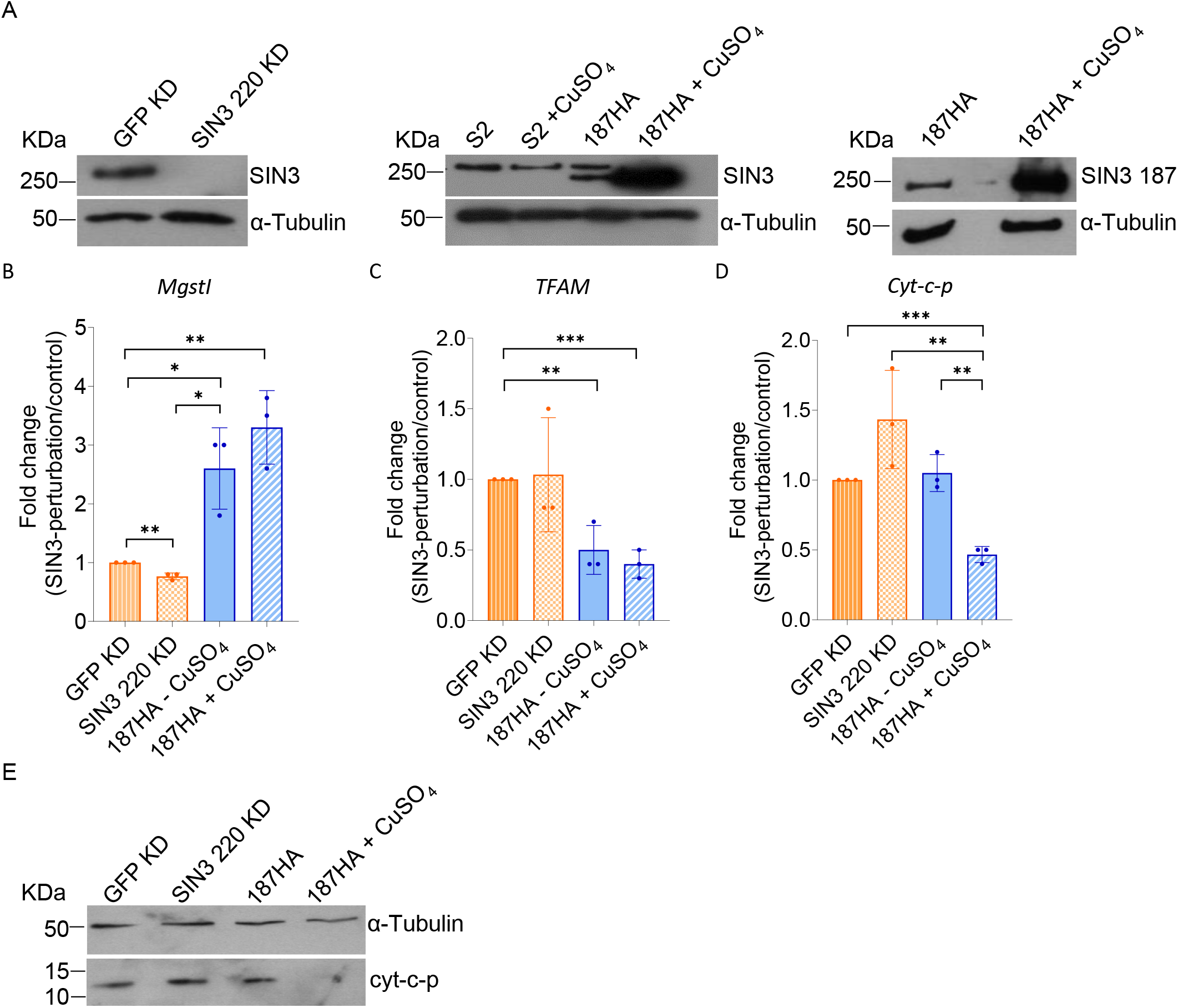
SIN3 187 regulates nuclear-encoded mitochondrial genes. *A,* Western blot analysis of whole cell extracts prepared from S2 and 187HA cells. Knockdown (KD) of *Sin3A* in S2 cells results in a decrease in the level of the dominant isoform, SIN3 220. Expression of HA-tagged SIN3 187 was induced via a metallothionine promoter by the addition of CuSO_4_. Blots were probed with SIN3 pan antibody that identifies all SIN3 isoforms or HA antibody to identify SIN3 187HA. α-Tubulin is used as the loading control. *B-D,* qRT-PCR analysis of *Mgst1 (B), TFAM (C),* and *Cyt-c-p (D).* Values indicated represent fold change for the specific SIN3 isoform perturbation, SIN3 220 knockdown or SIN3 187 overexpression, over GFP knockdown. Data are from a minimum of three biological replicates. Statistical comparisons between all conditions were done and significant changes are shown in the figure. Error bars represent standard error of the mean. *p*-value * *p*<0.05, ** *p*<0.01, *** *p*< 0.001. *E,* Western blot for cyt-c-p of whole cell extracts from indicated cells. α-Tubulin is used as the loading control.

We first validated the RNA-seq data for two nuclear-encoded mitochondrial targets that are bound by SIN3 187 (19). The first gene encodes microsomal glutathione S transferase (Mgst1), an enzyme that regulates response to ROS and influences longevity in fruit flies (22–23). We also selected mitochondrial transcription factor A (TFAM), known to influence mitochondrial DNA packaging, ATP production and generation of ROS (24–25). Our RNA-seq data indicate that the loss of SIN3 220 leads to reduced expression of *Mgst1* and does not affect the expression of *TFAM* (Table I). On the other hand, ectopic expression of SIN3 187 led to increased expression of *Mgst1* and reduced *TFAM* expression. We used quantitative real time PCR (qRT-PCR) to validate those findings and observed that the RNA-seq and qRT-PCR results are consistent. Indeed, *MgstI* expression was increased by ectopic expression of SIN3 187 and *TFAM* expression was repressed (Fig. 1*B, C*).

The RNA-seq data indicate that genes encoding several subunits from complex I, IV and V of the electron transport chain as well as *cytochrome c*, are repressed by SIN3 187. The data also show that SIN3 220 does not regulate the expression of genes encoding the listed subunits (Table I). The Drosophila genome has two *cyt c* genes. One of these genes codes for cyt-c-p, which is a part of the electron transport chain, and the other encodes cyt-c-d, which is mainly involved in the process of apoptosis (26). Our RNA-seq data reveal that *cyt-c-p* expression was repressed by SIN3 187. To confirm that finding, we analyzed gene and protein expression. Using qRT-PCR, we observed that the overexpression of SIN3 187 resulted in the repression of *cyt-c-p* expression (Fig. 1*D*). Additionally, in agreement with the gene expression data, the protein level of this subunit was strongly reduced with the overexpression of SIN3 187 (Fig. 1*E*).

Together, our results indicate that the ectopic expression of SIN3 187 negatively impacts expression of several nuclear-encoded mitochondrial genes. These targets include genes involved in oxidative phosphorylation, regulation of ROS and maintenance of mitochondrial function.

### SIN3 187 impacts mitochondrial respiration

During the process of mitochondrial respiration, the electron transport chain (ETC) subunits transport electrons via different complexes and pump protons across the mitochondrial membrane (27). The subunits ultimately transfer electrons to oxygen, resulting in the production of H_2_O. The observation that many ETC subunits are repressed by SIN3 187 led us to investigate whether mitochondrial respiration is affected under conditions in which we ectopically express this isoform. To address this, we used the Seahorse XFe96 system (Agilent), which can measure oxygen consumption and additional parameters pertaining to mitochondrial bioenergetics (28). Since Drosophila S2 cells express SIN3 220 as the dominant SIN3 isoform, we used these cells to obtain the respiration profile for SIN3 220. S2 cells treated with CuSO_4_ were used as an additional control in our experiments. Uninduced SIN3 187HA cells, with leaky expression of the small isoform, and SIN3 187HA cells treated with CuSO_4_ were used to obtain the respiration profile of SIN3 187 expressing cells. Treatment of S2 cells with CuSO_4_ did not appear to alter mitochondrial bioenergetics (Fig. 2). Compared to control S2 cells, the leaky and forced expression of SIN3 187 led to increased basal mitochondrial respiration (Fig. 2*B*). This result was surprising to us since we had expected that mitochondrial respiration would be reduced due to the altered expression of ETC subunits. Untreated and CuSO_4_ treated SIN3 187HA cells also had significantly higher spare respiratory capacity (SRC) compared to S2 cells (Fig. 2*C*). Interestingly, a study conducted in proliferative versus differentiated cells indicated that the SRC is higher in differentiated cells compared to their undifferentiated, proliferative counterparts (29). Previous work has established that SIN3 187 is the major isoform in differentiated tissues as well as in adult flies (17). Thus, cells expressing SIN3 187HA might be mimicking their natural contribution to regulating SRC.

**Figure 2.**
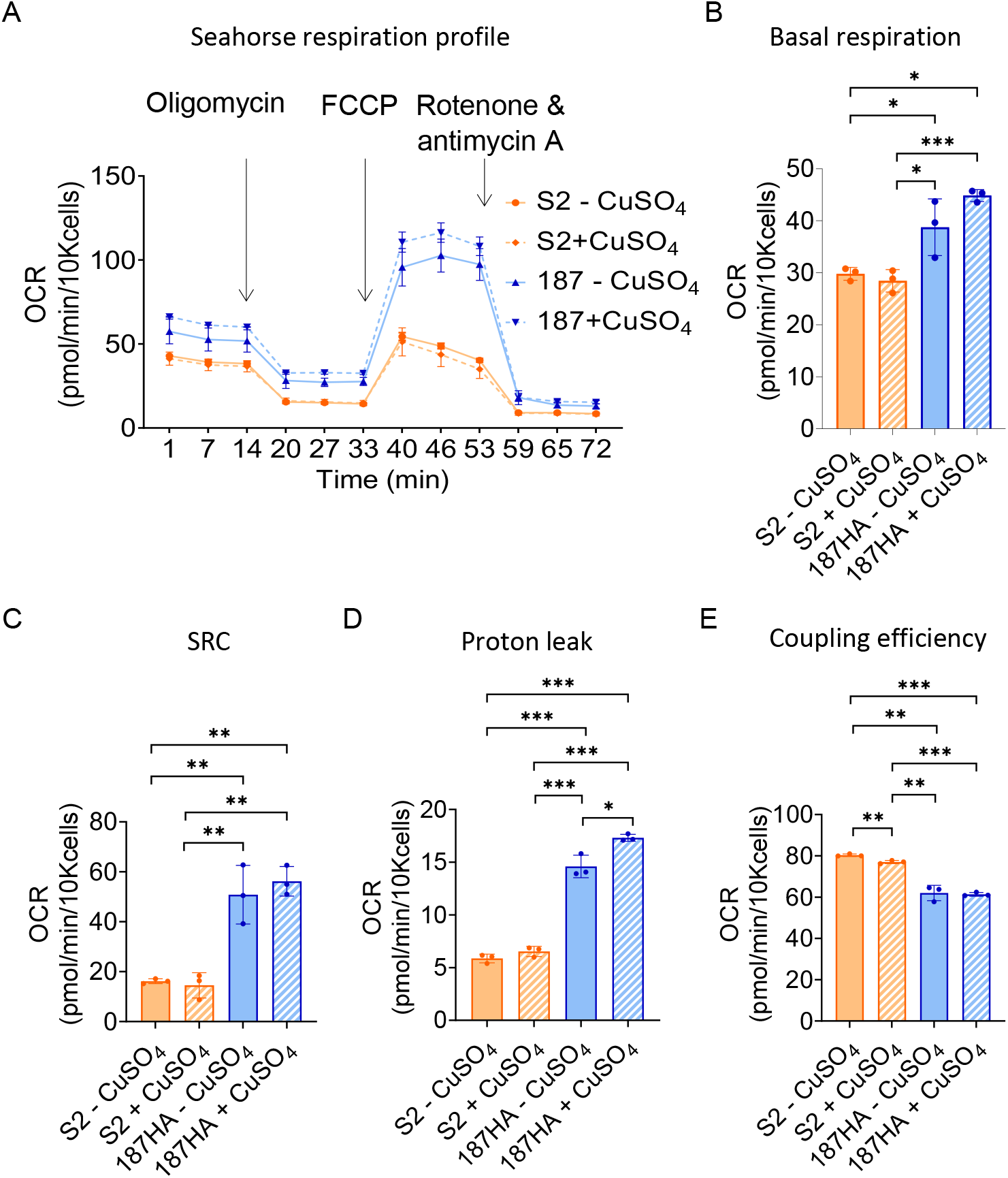
Ectopic expression of SIN3 187 affects mitochondrial respiration. *A,* Mitochondrial stress test profile of S2 and 187HA cells, each with and without CuSO_4_ treatment. Data are from three biological and eight technical replicates. *B-E*, Respiration profile from *(A)* was used to calculate Basal respiration *(B),* Spare respiratory capacity (SRC) *(C)*, Proton leak *(D),* and Coupling efficiency *(E).* Statistical comparisons between all conditions were done and significant changes are shown in the figure. Error bars represent standard error of the mean. *p*-value * *p*<0.05, ** *p*<0.01, *** *p*< 0.001.

The movement of protons across the mitochondrial membrane is typically linked to ATP production (29). Some protons can travel across the membrane without being coupled to ATP production, a phenomenon referred to as proton leak (29). The Seahorse respiration data can be used to determine the proton leak across the mitochondrial membrane. We determined that the leaky and forced expression of SIN3 187 led to increased proton leak, compared to the S2 cell counterparts (Fig. 2*D*). We speculate that the expression of SIN3 187 hinders optimal mitochondrial function and forces the cells to increase oxygen consumption and electron transport through the ETC in an attempt to regain homeostasis. Lastly, we calculated the overall coupling efficiency of cells expressing SIN3 220 versus those expressing SIN3 187. Coupling efficiency can be defined as the amount of mitochondrial respiration tied to ATP production (30). Cells expressing SIN3 187 had lower coupling efficiency compared to untreated and treated S2 cells (Fig. 2*E*). This finding indicates that SIN3 187 cells have elevated oxygen consumption but are unable to produce an equivalent energy output in terms of ATP production (Fig. 3*F*). Compared to younger tissues, older tissues have lower coupling efficiency (31). Since SIN3 187 is the dominant isoform in adult flies (17), reduced coupling efficiency might be mimicking the inherent property of this isoform. Together, our findings show that SIN3 187 regulates mitochondrial respiration. The repression of ETC subunits is correlated with negative impacts on proton leak and coupling efficiency. The ectopic expression of this isoform also results in higher SRC, which is characteristic of differentiated tissues.

**Figure 3.**
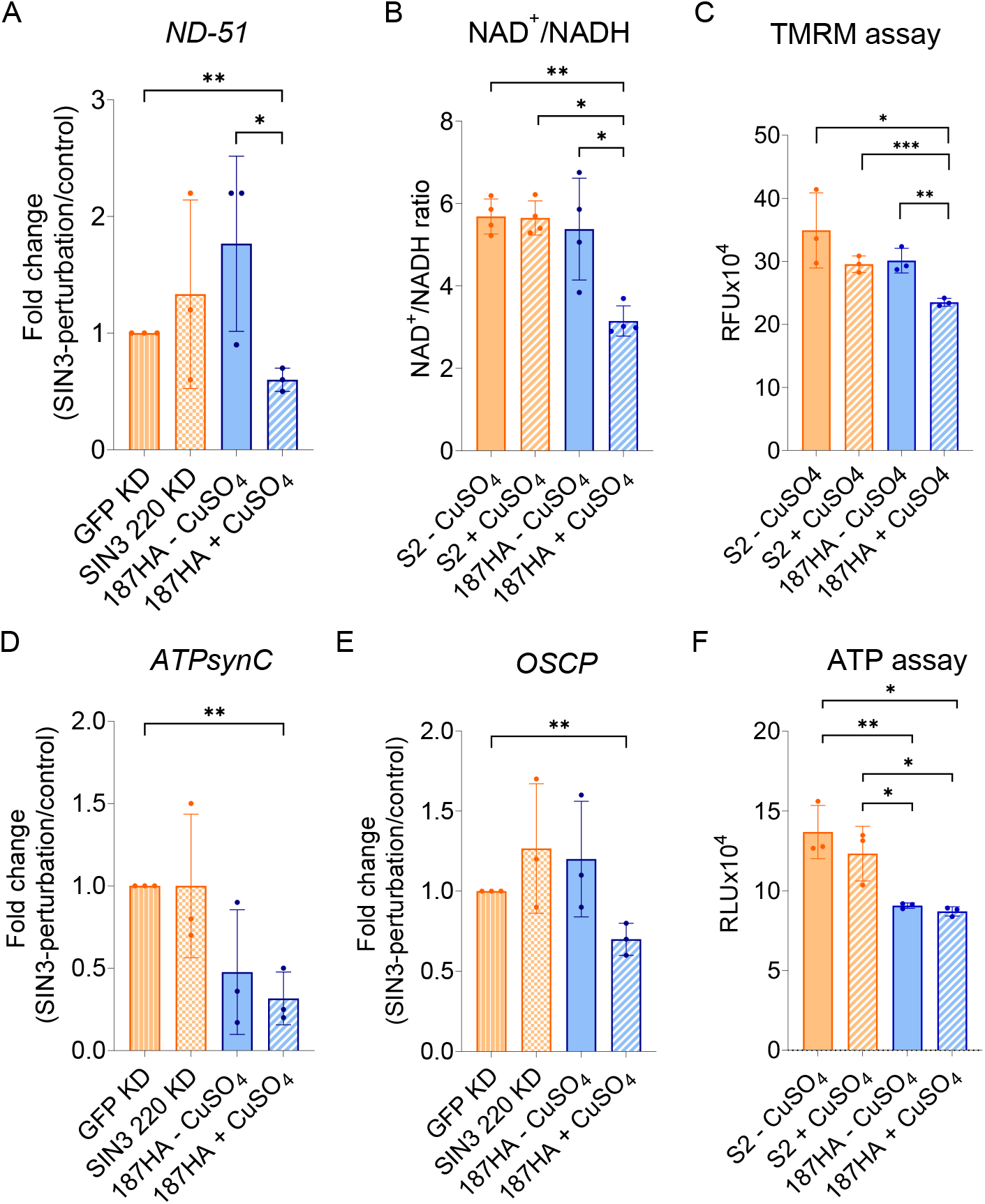
SIN3 187 affects mitochondrial bioenergetics. *A,* qRT-PCR used to measure expression of *ND-51* in the indicated samples*. B,* NAD^+^/NADH ratio in S2 and 187HA cells, with and without CuSO_4_. *C*, Mitochondrial membrane potential of S2 and 187HA cells estimated using TMRM, a fluorescent dye. Values shown are relative fluorescence units (RFU). *D-E*, qRT-PCR used to measure expression of *ATPsynC (D),* and *OSCP (E). F,* ATP levels of indicated cells as measured using a luminescence-based luciferase assay. Values shown in the graph are relative luminescence units (RLU). Data are from a minimum of three biological replicates. Statistical comparisons between all conditions were done and significant changes are shown in the figure. Error bars represent standard error of the mean. *p*-value * *p*<0.05, ** *p*<0.01, *** *p*< 0.001.

### SIN3 187 affects mitochondrial bioenergetics

ATP production is facilitated by electron transport and proton pumping through the five complexes of the ETC (27, 32). The process starts with the transfer of electrons from NADH to complex I, resulting in the production of NAD^+^ (32). Cells maintain an optimal NAD^+^/NADH ratio, which is crucial for mitochondrial homeostasis (33). Disruption of this ratio is indicative of mitochondrial ROS production (27). In mammalian cells, deficiencies in mitochondrial subunits of complex I affect oxidative phosphorylation and the NAD^+^/NADH ratio (34). Complex I is the largest ETC complex, comprised of several subunits (35). The NDUFV1 subunit of this complex acts as the NADH acceptor site.

Our analysis of the published RNA-seq data indicates that the expression of the Drosophila ortholog of *NDUFV1*, known as *ND-51*, is repressed by SIN3 187 (19) (Table I). To confirm this finding, we used qRT-PCR analysis and noted that indeed the overexpression of SIN3 187 led to a reduction in expression of *ND-51* (Fig. 3*A*). Next, we sought to investigate whether the NAD^+^/NADH ratio was affected by measuring the levels of these metabolic intermediates and their relative ratio. Treatment of S2 cells with CuSO_4_ did not affect the NAD^+^/NADH ratio (Fig. 3*B*). Similarly, the leaky expression of the SIN3 187 isoform did not alter the ratio of the metabolic intermediates. The cells forced to overexpress SIN3 187, however, showed a significant reduction in the ratio, suggesting that relative level of the isoforms is important in control of the ratio (Fig. 3*B*).

As electrons are transported through the ETC complexes, protons are pumped from the mitochondrial matrix to the intermembrane space (36). This action creates a concentration difference across the membrane, which is used by complex V for ATP production. Since the ectopic expression of SIN3 187 led to repression of several ETC subunits, we asked whether the pumping of protons, and thereby the membrane potential, would be affected. To investigate this possibility, we used the TMRM dye, a positively charged dye that accumulates in the negatively charged mitochondrial matrix (37). Failure to maintain a potential difference across the membrane results in loss of negative charge in the matrix and low accumulation of the dye. The mitochondrial membrane potential did not change with the CuSO_4_ treatment of S2 cells. Similarly, the leaky expression of SIN3 187 did not significantly affect the membrane potential. Overexpression of SIN3 187, however, resulted in a significant reduction in membrane potential (Fig. 3*C*). This finding suggests that the repression of several ETC subunits negatively affects proton pumping and maintenance of a potential charge difference across the membrane in the presence of elevated levels of SIN3 187.

Lastly, we wanted to study the effect of ectopic expression of SIN3 187 on ATP production. The potential difference generated by the pumping of protons across the mitochondrial membrane is harnessed by complex V (ATP synthase) to produce ATP (36). The ATP synthase complex is composed of several subunits that are organized into the F_o_-F_1_ domains, connected by a peripheral stalk. Published RNA-seq data indicate that several complex V subunits are repressed by SIN3 187 (19) (Table I). Two of these candidate genes, *ATPsynC* and *OSCP*, are known to affect ATP production (38–39). Using qRT-PCR, we confirmed that the expression of *ATPsynC* and *OSCP* are repressed by the forced expression of SIN3 187 (Fig. 3, *D* and *E*). Given these data, we hypothesized that the disruption of the mitochondrial membrane potential, combined with the repression of ETC subunits, would lead to a reduction in ATP synthesis. Using a luminescence-based assay, we determined the total amount of ATP in our S2 and 187HA cells. As predicted, we observed a significant reduction in ATP levels in cells induced to overexpress SIN3 187 (Fig. 3*F*). We observed that the uninduced SIN3 187HA cells also had a similarly low level of ATP (Fig. 3*F*). This might be due to the leaky expression of SIN3 187 in these cells (Fig. 1*B*). Overall, these data show that SIN3 187 represses the expression of several nuclear-encoded mitochondrial genes. This altered gene expression profile is associated with disruption of mitochondrial function, which is manifested as changes in proton leak, mitochondrial membrane potential and ATP production.

### SIN3 187 affects ROS production

ROS are inevitably produced as electrons are transferred through the subunits of the ETC (40). During the process of mitochondrial respiration, some electrons are transferred to oxygen molecules, resulting in the formation of ROS (40). Complexes I, II and III are the main sites of ROS production (27, 40). Two separate studies using fibroblast cells derived from patients with low complex I activity demonstrated that the reduced activity of the complex results in higher ROS production (41–42). Based on the effects of forced expression of SIN3 187 on mitochondrial bioenergetics, we hypothesized that these cells would have high ROS levels. Using a luminescence assay, we measured ROS in cells expressing SIN3 220 versus those expressing SIN3 187. Treatment of S2 cells with CuSO_4_ did not affect ROS levels (Fig. 4*A*). The leaky expression of SIN3 187 in untreated SIN3 187 cells, however, resulted in a small yet statistically significant increase in ROS production. This level further increased when we forced the expression of the SIN3 187 isoform using CuSO_4_ (Fig. 4*A*). These results demonstrate that the ectopic expression of SIN3 187 affects ROS levels in Drosophila.

**Figure 4.**
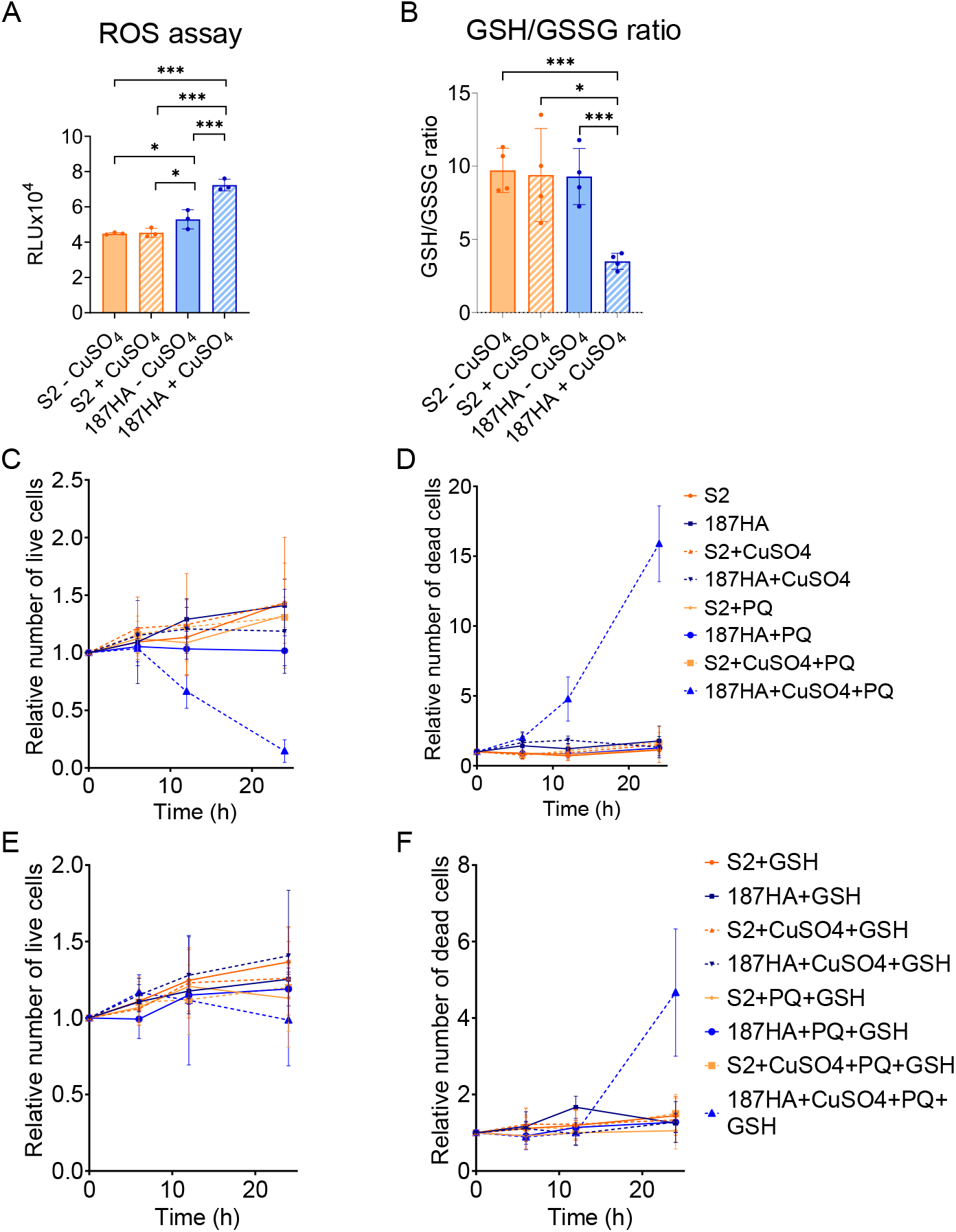
SIN3 187 affects ROS production. *A*, ROS production measured using a luminescence-based assay in S2 and 187HA cells, with and without CuSO_4_. Values shown are relative luminescence units (RLU). *B,* GSH/ GSSG ratio quantified using a luminescence-based kit in indicated S2 and 187HA cells. *C-D,* Quantification of live *(C),* and dead cells *(D)* with indicated treatments. *E-F,* Quantification of live *(E),* and dead *(F)* S2 and 187HA cells treated with GSH. Cell viability determined by the trypan blue staining method. The relative number of cells was calculated using the formula: (number of live or dead cells at time point X/live or dead cells at 0 hr time point)/ total number of cells for time point X. Data are from a minimum of three biological replicates. Statistical comparisons between all conditions were done and significant changes are shown in the figure. Error bars represent standard error of the mean. *p*-value * *p*<0.05, ** *p*<0.01, *** *p*< 0.001.

Reduced glutathione (GSH) acts as a scavenger to remove ROS intermediates (43). The ratio of GSH to its oxidized form, GSSG, is thus relatively high during low oxidative stress conditions (43–44). Based on the mitochondrial dysfunction phenotype observed, we measured the GSH/GSSG ratio in our cells. The treatment of S2 cells with the CuSO_4_ inducer did not alter this ratio compared to untreated S2 cells (Fig. 4*B*). Similarly, the leaky expression of SIN3 187 did not affect the GSH/GSSG ratio. The forced expression of SIN3 187, however, resulted in a significant reduction of the ratio, indicative of increased ROS accumulation (Fig. 4*B*). These data indicate that overexpressing SIN3 187 perturbs the ROS levels in the cell. This could be attributed to the sub-optimal functioning of multiple ETC subunits.

We hypothesized that since cells forced to express SIN3 187 exhibited mitochondrial dysfunction combined with altered ROS levels, they might show increased sensitivity to an external source of ROS. To evaluate the sensitivity of cells expressing SIN3 187 to oxidative stress, we used paraquat, a well-known stressor (45). We assessed viability of S2 versus SIN3 187 cells over a period of 24 hours, with or without the addition of paraquat. Untreated and CuSO_4_ treated S2 and SIN3 187 cells did not demonstrate any proliferation defect in this time period (Fig. 4, *C* and *D*). For all samples, we observed a small increase in the number of live cells, with no large changes in the number of dead cells relative to the starting cell cultures. When we treated S2 and SIN3 187 cells with paraquat, we did not detect any changes in the number of dead cells. There was a small but insignificant reduction in the number of live cells, indicating that the concentration of paraquat we used was non-toxic to the cells. S2 cells treated with both CuSO_4_ and paraquat remained viable and divided at a similar rate compared to non-treated control cells. Dual treatment of SIN3 187 cells with CuSO_4_ and paraquat, however, resulted in a large reduction in the number of live cells. This difference in viability was apparent 12 hours post treatment with paraquat. The number of dead cells increased at the same time, showing that these cells have a higher susceptibility to paraquat toxicity compared to controls. Comparing the 24-hour readings of untreated SIN3 187 cells to dual treated SIN3 187 cells, we observed a 9-fold decrease in the number of live cells and an 8.9-fold increase in the number of dead cells. These results indicate that cells forced to predominantly express SIN3 187 are more sensitive to external oxidative stress compared to cells expressing SIN3 220. This sensitivity might stem from the low GSH/GSSG ratio, indicating a reduced ability to neutralize ROS.

Supplementation of cells with the reduced form of glutathione (GSH) can rescue ROS induced toxicity (46–47). Here, we hypothesized that the addition of reduced glutathione (GSH) would rescue the observed sensitivity to paraquat in cells forced to express SIN3 187. We pre-treated our cells twice with GSH, once before addition of CuSO_4_ and again before addition of paraquat. Sole treatment with GSH did not affect viability and proliferation of S2 and SIN3 187 cells (Fig. 4, *E* and *F*). Similarly, we did not observe any changes in the relative number of live and dead cells when samples were treated with GSH and paraquat. S2 cells that were treated with CuSO_4_, paraquat and GSH maintained a similar viability profile to those without GSH treatment. The GSH treatment of SIN3 187 cells exposed to CuSO_4_ and paraquat resulted in over a 6-fold improvement in the number of live cells at the end of 24-hour treatment with paraquat. We also observed over a 3-fold reduction in the number of dead cells. This finding confirmed that cells forced to express SIN3 187 are sensitive to external oxidative stress, which can be largely rescued by glutathione supplementation.

### SIN3 187 regulates the apoptotic pathway

Overproduction of ROS from dysfunctional mitochondrial can result in apoptosis (48). Based on our previous publications, we noted that the overexpression of SIN3 187 results in the activation of several pro-apoptotic genes while repressing a few apoptotic inhibitors (Table II) (12, 19). This regulation was unique to SIN3 187.

**Table II:**
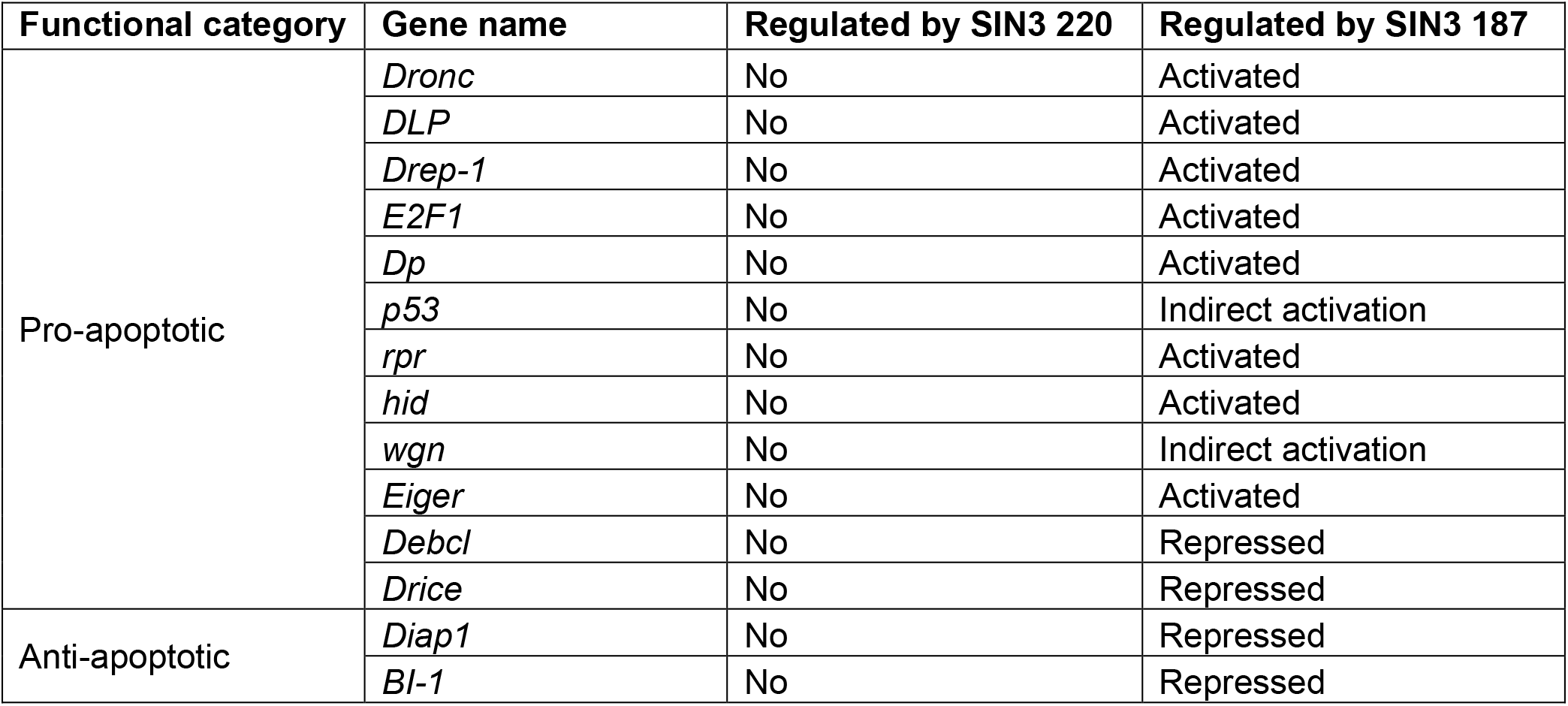
Nuclear-encoded apoptotic genes are differentially regulated by SIN3 isoforms. Genes that are both bound by SIN3 and altered in expression following perturbation in the level of the specific isoform are considered direct targets. The direction of regulation (activated or repressed) is indicated. Indirectly regulated genes are those that change in expression with the perturbation of the SIN3 isoform but the isoform was not found to bind at the promoter. Data obtained by analyzing published data sets (12, 19).

| Functional category | Gene name | Regulated by SIN3 220 | Regulated by SIN3 187 |
| --- | --- | --- | --- |
| Pro-apoptotic | *Dronc* | No | Activated |
|  | *DLP* | No | Activated |
|  | *Drep-1* | No | Activated |
|  | *E2F1* | No | Activated |
|  | *Dp* | No | Activated |
|  | *p53* | No | Indirect activation |
|  | *rpr* | No | Activated |
|  | *hid* | No | Activated |
|  | *wgn* | No | Indirect activation |
|  | *Eiger* | No | Activated |
|  | *Debcl* | No | Repressed |
|  | *Drice* | No | Repressed |
| Anti-apoptotic | *Diap1* | No | Repressed |
|  | *Bl-1* | No | Repressed |

To validate our RNA-seq data, we selected two pro-apoptotic genes, *Dronc*, the caspase 9 homolog in Drosophila, and *p53*, a well-known activator of the apoptotic pathway. Dronc is an initiator caspase that when cleaved and activated, forms an apoptosome complex to initiate the apoptotic pathway (49–50). p53 is a transcription factor that modulates the apoptotic response by affecting transcription of apoptosis related genes (51). Apart from the transcriptional regulation of apoptosis, p53 is capable of inducing apoptosis via non-transcriptional mechanisms (51). Our qRT-PCR data show that loss of SIN3 220 by RNAi does not affect the expression of *Dronc* or *p53* (Fig. 5, *A* and *B*). The leaky expression of SIN3 187 was sufficient to lead to upregulation of *Dronc*, while the overexpression of this isoform resulted in significant upregulation of both pro-apoptotic genes tested. These findings indicate that SIN3 187 influences the expression of genes in the apoptotic pathway in Drosophila.

**Figure 5.**
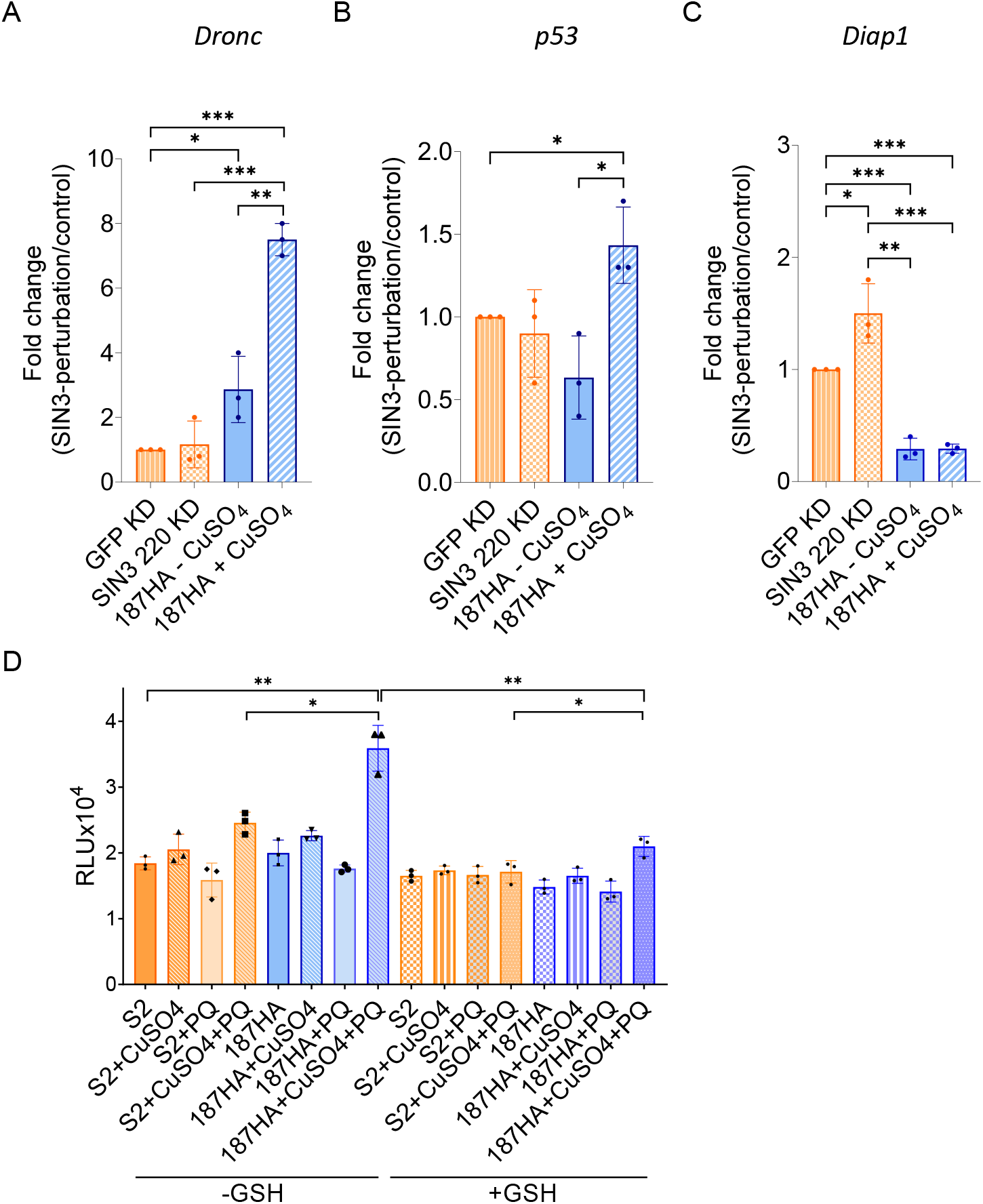
SIN3 187 regulates the apoptotic pathway. *A-C,* qRT-PCR used to measure expression of apoptotic pathway genes *Dronc (A), p53 (B),* and *Diap1 (C).* Statistical comparisons between all conditions were done and significant changes are shown in the figure. *D,* Caspase assay showing the levels of active caspases in all cell conditions tested, with and without GSH. Statistical comparisons between all conditions were done and limited statistical comparisons are shown in the figure. Data are from a minimum of three biological replicates. Error bars represent standard error of the mean. *p*-value * *p*<0.05, ** *p*<0.01, *** *p*< 0.001.

The activation of Dronc is inhibited by its binding to the apoptotic inhibitor Diap1 (49). Loss of Diap1 results in the activation of procaspase Dronc into its catalytically active form (52). Using qRT-PCR, we observed that the leaky expression of SIN3 187 was enough to repress the expression of *Diap1* and this was accompanied by an upregulation in the expression of *Dronc* (Fig. 5, *B* and *C*). The overexpression of SIN3 187 also results in the repression of *Diap1*, however, this level is comparable to that of uninduced SIN3 187 cells. Overall, our data suggest that the ectopic expression of SIN3 187 leads to the repression of a few anti-apoptotic and activation of several pro-apoptotic genes. These genes work at different levels in the apoptotic pathway, indicating that SIN3 187 likely impacts the execution of the apoptotic pathway at various stages.

Lastly, to confirm the loss of cell viability by execution of the apoptotic pathway, we investigated the levels of active caspases. Using the same set of cells used to test oxidative stress susceptibility, we measured the levels of active caspases. Cells forced to ectopically express SIN3 187 showed a small upregulation of active caspases (Fig. 5*D*). Based on our cell proliferation data, this upregulation is not enough to result in cell death. This result indicates that ectopic overexpression of this isoform does not by itself lead to lethality by apoptosis. SIN3 187 expressing cells treated with paraquat showed the highest increase in levels of active caspases. The activation of caspases was rescued by treatment of cells with GSH (Fig. 5*D*). Treatment of SIN3 187 expressing cells with paraquat and GSH showed a significant reduction in levels of active caspases when compared to cells not treated with the antioxidant. Taken together, these results suggest that the forced expression of SIN3 187 activates the expression of pro-apoptotic genes. This equips the cells with increased ability to execute cell death via apoptosis in the presence of external oxidative stress. Addition of glutathione neutralizes ROS levels and suppresses cell death by apoptosis.

## Discussion

As a transcriptional regulator, SIN3 is known to regulate genes involved in a variety of cellular pathways, ranging from those controlling cell proliferation and metabolism to apoptosis (19, 53). In this paper, we present a novel comparison of the roles of Drosophila SIN3 isoforms, SIN3 220 and SIN3 187, in the regulation of two essential processes, mitochondrial respiration and response to oxidative stress. For these studies, we utilized cell lines that predominantly express either SIN3 220 or SIN3 187 (Fig. 1*A*). This cultured cell system enabled us to study the role of each of the isoforms in regulating the pathway of interest.

Previously, we have seen that SIN3 220 regulates the expression of many nuclear-encoded mitochondrial genes. Using published RNA-seq data, we identified targets of SIN3 187 that are crucial to the process of oxidative phosphorylation (19). Interestingly, a large majority of these genes were not regulated by SIN3 220. Several of these genes encode subunits of the ETC (Table I). Other identified genes are responsible for maintenance of various aspects of mitochondrial function. One such target, MgstI, plays a role in regulating response to oxidative stress and cell death (22). A site on the Mgst1 protein can act as a sensor to recognize ROS intermediates. Another target of SIN3 187 is TFAM. In mice, TFAM modulates the expression of mitochondrial DNA (mtDNA) encoded genes, mtDNA replication as well as its maintenance (24, 54). Mice expressing a mutated version of TFAM show sub-optimal function of several electron transport chain subunits in the mitochondria. Here, we observed that ectopic expression SIN3 187 results in the upregulation of oxygen consumption, increases proton leak and lowers coupling efficiency (Fig. 2). We postulate that repression of ETC subunits leads to their sub-optimal function. This might lead the cells to upregulate oxygen consumption and transport through the ETC to compensate for reduced efficiency of the ETC. A study comparing young versus aged rat cardiomyocytes demonstrated that older cells have a comparatively higher basal respiration rate that can be ascribed to their increased proton leak (55). A similar mechanism may explain our current findings.

The repression of ETC subunits also affects other parameters associated with mitochondrial respiration. The repression of complex I subunits in mammalian cell lines affects the NAD^+^/NADH ratio (56). Yeast strains carrying mutant versions of a subunit of complex I, NDUFV1, are unable to oxidize NADH to NAD^+^ (57). Similarly, knockdown of this subunit in breast cancer cells reduces the NAD^+^/NADH ratio (58). Consistent with those studies in other organisms, we show here that SIN3 187 repressed *ND-51*, the homolog of *NDUFV1*, and the cells correspondingly have a reduced NAD+/NADH ratio as compared to S2 cells (Fig. 3, *A* and *B*).

SIN3 187 also represses some complex V subunits, such as *ATPsynC* and *OSCP*. ATPsynC is a part of the F_1_ domain of complex V. Drosophila strains expressing mutant versions of ATPsynC have reduced production of ATP along with diminished locomotor function (38). OSCP is a nuclear-encoded mitochondrial subunit localized to the stalk of complex V. Knockdown of *OSCP* in mouse neuronal cells results in a reduction of mitochondrial membrane potential and ATP production (39). These studies underscore the importance of complex V in energy production. We demonstrate that ectopic expression of SIN3 187 led to the repression of complex V subunits (Fig. 3, *D* and *E*). The lowered expression of these subunits along with reduced mitochondrial membrane potential may cause reduced ATP production in cells overexpressing SIN3 187 (Fig. 3*F*).

Disruption of mitochondrial membrane potential together with ETC dysfunction and a decrease in ATP production are all indicative of oxidative stress (48, 59). We find that cells forced to express SIN3 187 have higher ROS production and a lower GSH/GSSG ratio compared to those expressing SIN3 220 (Fig. 4, *A* and *B*). Using paraquat as a source of ROS, we observe that the forced expression of SIN3 187 greatly increases sensitivity to oxidative stress (Fig. 4, *C* and *D*). This finding could be explained by the inversed GSH/GSSG ratio, which could diminish the ability of these cells to neutralize ROS. Rescue of cell viability by addition of GSH provides further evidence to support this hypothesis (Fig. 4, *E* and *F*).

Additional investigation into the GO categories influenced by SIN3 187 unveiled that of the two SIN3 isoforms, SIN3 187 exclusively regulates the expression of apoptotic genes (Table II). Upregulation of pro-apoptotic and repression of anti-apoptotic genes might poise cells expressing SIN3 187 to the execution of the apoptotic pathway. When faced with an external source of oxidative stress, cells expressing SIN3 187 can readily initiate the apoptotic cascade, manifested as increased sensitivity. The process of aging is accompanied by changes in cellular physiology. Based on the “mitochondrial theory of aging,” decline in mitochondrial function and increased ROS are commonly observed during aging (60–62). High levels of ROS in aging tissues are thought to be a contributor to the progressive decline in cellular homeostasis (61). ETC dysfunction combined with changes to ATP have been noted in aging tissues (63). Since SIN3 187 is the dominant SIN3 isoform in differentiated cells and in adult flies (17), we predict that this isoform might contribute to the process of aging. Additionally, since SIN3 187 regulates genes in the apoptotic pathway, it might be instrumental in the removal of aged and damaged cells. Clearing the organism of damaged and senescent cells could aid in maintenance of homeostasis.

Interestingly, some of the phenotypes observed from cells exhibiting leaky expression of the SIN3 187HA transgene were similar to those obtained using cells forced to overexpress the transgene. For example, we found no significant difference in basal respiration and ATP levels between the two cell types (Fig 2*B*, 3*F*). Similarly, the expression of select pro-apoptotic genes showed the same trend in gene expression between the leaky and forced expression of SIN3 187 (Fig 5*A* and *C*). These data suggest that in cells that express both SIN3 isoforms, SIN3 187 is able to outcompete the action of SIN3 220 to influence cellular physiology in a manner similar to cells that predominantly express SIN3 187. To investigate this idea further, future studies will include use of tissues and developmental stages where each isoform is naturally predominant.

In summary, we find that two SIN3 isoforms, SIN3 220 and SIN3 187, differentially regulate genes essential for the maintenance of energy metabolism and stress response. Given that SIN3 isoforms are differentially expressed during development (17), the predominant complex and its regulated target genes will dictate a specific response appropriate to the needs of that given cell or tissue. We speculate that the presence of unique SIN3 isoform interacting partners might be instrumental in differentially regulating target genes and controlling the cell response. For example, since the demethylase dKDM5/LID preferentially interacts with SIN3 220 (18), SIN3 220 complexes may impact both histone acetylation and methylation at gene targets, while we predict that SIN3 187 only affects acetylation. Future studies will be aimed at investigating the mechanisms through which distinct SIN3 isoform complexes regulate cellular homeostasis.

## Experimental procedures

### Maintenance of cell culture

Drosophila Schneider 2 (S2) lines were maintained in 1 X Schneider’s medium supplemented with 10% heat inactivated fetal bovine serum (Gibco) and 50 mg/ml gentamicin antibiotic (Gibco). Transgenic S2 cells carrying the SIN3 187 transgene tagged with an HA epitope were grown in medium containing 0.1 mg/ml penicillin and streptomycin (Gibco). For treatment with CuSO_4_, cells were incubated with 0.035 M of the inducer for 36 hr. All Drosophila cell culture lines were maintained at 27°C.

### RNAi protocol

The RNAi protocol followed was the same as described (64). Western blotting was used to confirm knockdown of *Sin3A*.

### Western blot analysis

Western blotting was performed as described (64). In brief, 12 ug of whole cell protein was used for all western blots probing for SIN3 and tubulin. 50 ug of protein was loaded while probing for cyt-c-p. Protein concentrations were determined using the Protein DC assay, following the manufacturer’s protocol (Bio-Rad). Proteins were separated on 8% SDS gels and transferred to PVDF membranes (Thermo Fisher Scientific). A 15% SDS gel was used to separate proteins for detection of cyt-c-p. Membranes were incubated with 5% blocking milk solution for 1 hr, followed by washing with 1 X PBS buffer with 0.2% Tween 20 (PBST). Blots were incubated with the primary antibody for 2 hr, unless otherwise indicated. All blots were incubated with secondary antibody (1:3000) for 1 hr at room temperature and then signals detected using ECL Prime (Cytiva Life Sciences). Primary antibodies used were SIN3 (1:1000) (65), α-tubulin (1:1000, incubated overnight in 5% BSA in 1 X PBST at 4°C, Cell Signaling), HA-HRP (1:6000, Sigma), cyt-c-p (1:100, incubated overnight at 4°C, Thermo Fisher Scientific). All biological replicates were tested by western blotting.

### Real-time qRT-PCR

Complementary DNA was prepared from total RNA extracted from indicated samples using the ImProm-II Reverse Transcription System (Promega). qRT-PCR analysis was performed using a master mix with ROX reference dye (Invitrogen), SYBR Green I nucleic acid gel stain and Go Taq Hot Start Polymerase (Promega). qRT-PCRs were carried out in a QuantStudio 3 Real-Time PCR system (Thermo Fisher Scientific). Actin was used as an internal control to normalize levels of RNA. Primers used for amplification are listed in Supplementary Table I.

### Seahorse respiration assay

The Agilent Seahorse XFe96 Analyzer was used to measure oxygen consumption in intact cells, with the Seahorse XF Cell Mito Stress Test Kit (103015-100). Using the oxygen consumption output, various parameters of mitochondrial respiration including basal respiration, proton leak, spare respiratory capacity and coupling efficiency were measured using formulas specified by the manufacturer. Pharmacological agents from the kit used for the study were Oligomycin (2 μM), FCCP (0.75 μM), Rotenone and Antimycin A (0.5 μM). The experiment was performed with eight technical and three biological replicates for each cell type tested. All values were normalized to a cell number of 10,000.

### Determining NAD^+^/NADH ratio and GSH/GSSG ratio

Total cellular NAD+/NADH and GSH/GSSG ratios were determined using luminescence based kits (Promega). Manufacturer’s protocols were used without any deviations. Luminescence was read using the SpectraMax i3x plate reader (Molecular Devices). The experiments were performed with a minimum of two technical and three biological replicates for each cell type tested. All values were normalized to cell number.

### TMRM assay

Mitochondrial membrane potential was determined using 20 mM Tetramethylrhodamine (TMRM) dye (Invitrogen, I34361, T668). Briefly, 100,000 cells were added per well and treated with TMRM for 30 minutes. The plate was centrifuged, and the cells were washed with 1 X PBS. Fluorescence was read using the SpectraMax i3x plate reader (Molecular Devices). The experiment was performed with 12 technical and three biological replicates for each cell type tested. All values were normalized to cell number.

### Cellular ATP levels

Intracellular levels of ATP were determined using the CellTitre-Glo 2.0 Cell Viability assay (Promega). Cells were treated with the CuSO_4_ inducer for 36 hours before measuring ATP. 10,000 cells per treatment condition were used to assess ATP levels, following the manufacturer’s protocol. Luminescence was read using the SpectraMax i3x plate reader (Molecular Devices). The experiment was performed with six technical and three biological replicates for each cell type tested. All values were normalized to cell number.

### ROS assay

Cellular ROS levels was detected using the ROS-Glo H_2_O_2_ assay as per manufacturer’s instructions (Promega). Cells were treated with CuSO_4_ for 36 hr before carrying out the assay. Luminescence was read using the SpectraMax i3x plate reader (Molecular Devices). The experiment was performed with two technical and three biological replicates for each cell type tested. All values were normalized to cell number.

### Cell proliferation assay

For all experiments, cells were counted using trypan blue stain (Lonza Bioscience). For the paraquat proliferation assays, cells were treated with 0.035 M CuSO_4_ for 36 hr. Following the incubation, cells were treated with 10 mM paraquat (Sigma-Aldrich). Using the trypan blue stain, live and dead cells were counted at indicated time points over a period of 24 hr. For the proliferation assay with paraquat and GSH, cells were treated with 5 mM L-Glutathione reduced (Sigma-Aldrich) for 1 hr before adding CuSO_4_ inducer. After this 36 hr induction period, 5 mM of L-Glutathione reduced was added again and the cells were incubated for 1 hr before the addition of 10 mM paraquat (Sigma-Aldrich). Cells were then counted at indicated time points over a period of 24 hr using trypan blue stain.

### Apoptosis assay

Active caspases were detected using the Caspase-Glo 3/7 Assay System (Promega). The experiment was set up using the same protocol as proliferation assay. Cells were harvested 8 hr after treatment with paraquat and tested for active caspases. Luminescence was read using the SpectraMax i3x plate reader (Molecular Devices). The experiment was performed with three biological replicates for each cell type tested. All values were normalized to cell number.

## Data availability

All data generated for this study are contained within the manuscript and can be shared upon request.

## Supporting information

This article contains supporting information.

## Acknowledgements

We thank members from the laboratories of Dr. Pile and Dr. Greenberg for helpful discussions and suggestions throughout the time of conducting this research.

## Author contributions

A.M., A.C., and L.A.P. conceptualization; A.M., L.V., I.S., investigation; A.M., and L.A.P. writing-original draft; A.M., L.V., I.S., A.C., M.G., and L.A.P. review and editing; M.L.G., and L.A.P. supervision.

## Funding and additional information

This work was supported by Office of the Vice President for Research, Wayne State University to L.A.P., as well as National Institutes of Health grant numbers R01GM125082 and R01HL117880 to M.L.G.

## Abbreviations

ChIP-seq: ChIP sequencing
CuSO_4_: copper sulfate
ETC: electron transport chain
ROS: reactive oxygen species
GSH: reduced glutathione
GSSG: glutathione disulfide
HDAC: histone deacetylase
MEF: murine embryonic fibroblasts
SRC: spare respiratory capacity
qRT-PCR: quantitative RT-PCR

