## Supplemental Table I for "Isoforms of the transcriptional cofactor SIN3 differentially regulate genes necessary for energy metabolism and cell survival"

**Supplementary Table I**:

Primers used for qRT-PCR

| **Primer** | **Sequence** |
| --- | --- |
| *MgstI* forward | CCCAAGCTGAAGGT AAG TT |
| *MgstI* reverse | CGGATCAGTCAGGACGTAGA |
| *TFAM* forward | AAGCGGGTCAAGGAGCTG |
| *TFAM* reverse | GGAAATCGCTTTCCTGTAGAG C |
| *Cyt-c-p* forward | GCTCGACGTTTGTGTTCAAT |
| *Cyt-c-p* reverse | TTCCCTTCTCAACATCACCA |
| *ND-51* forward | TTGGTGGTGAATGCCGATGA |
| *ND-51* reverse | AGCCTCGTTGTAGAACTCGC |
| *ATPsyn C* forward | GCCGCAACAGTCGGTGTC |
| *ATPsyn C* reverse | AGGCGAACAGCAGCAGGAA |
| *OSCP* forward | TACAAGACCATCATGGCCGC |
| *OSCP* reverse | ATCAGGCCACCAATGATGCT |
| *Dronc* forward | TGAGTCCGATTCAAGGCCAC |
| *Dronc* reverse | CCGTCAACGACACCCACATA |
| *p53* forward | CCTTCCCCAACAAGATCGCT |
| *p53* reverse | ACCTCCACCGTTTTCGGAAT |
| *Diap1* forward | GTCAAATCTCAACGCAACGGA |
| *Diap1* reverse | GAAGGTAACCGCAGAGGTCC |
| *Actin* forward | CGCAGCTCATTGTAGAAGGT |
| *Actin* reverse | CTGGGACGATATGGAGAAGA |
