## Supplementary figures and images for "Isoforms of the transcriptional cofactor SIN3 differentially regulate genes necessary for energy metabolism and cell survival"

### Supplemental figure 1

#### Slide 1
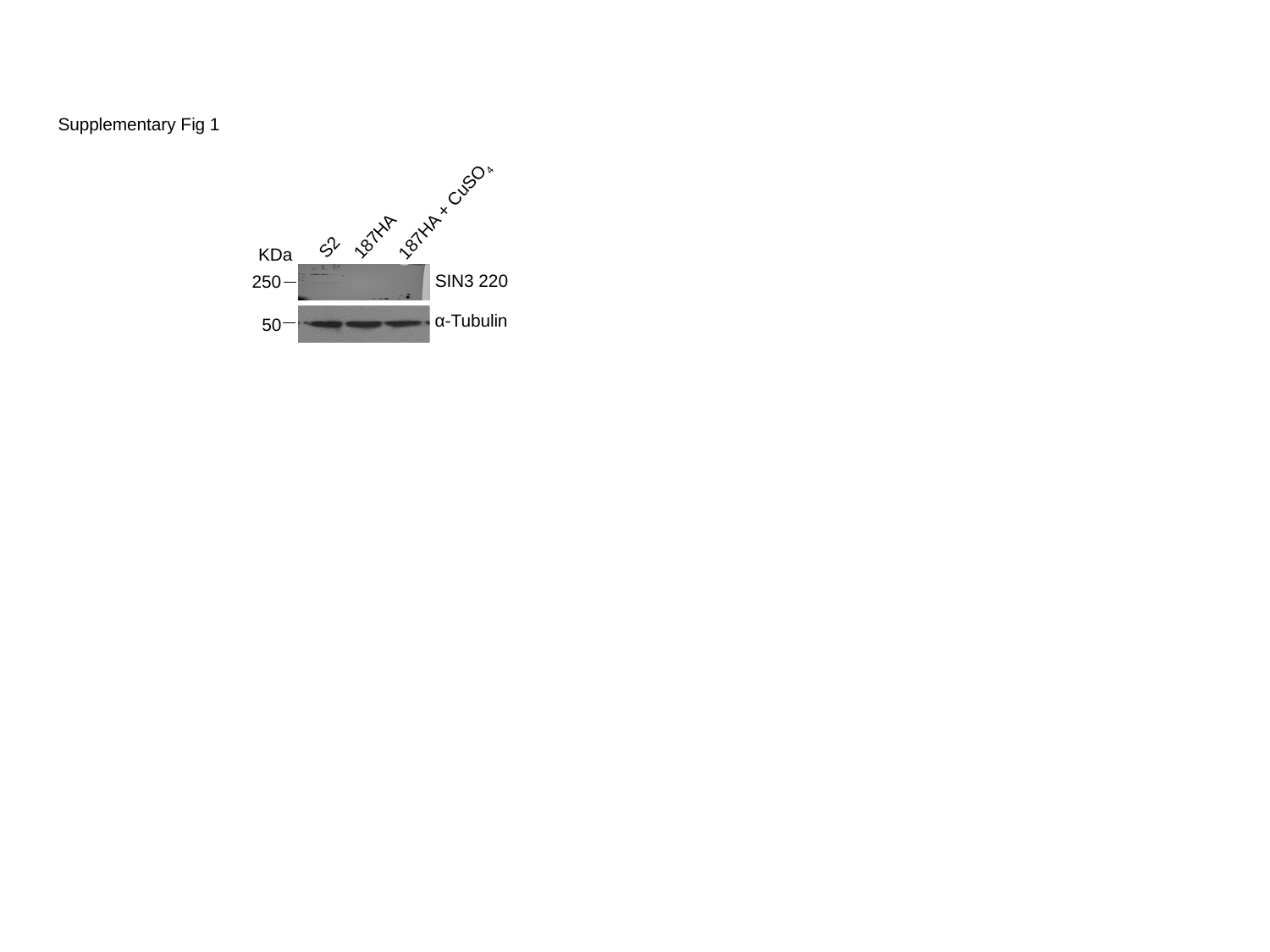

187HA + CuSO_4_
187HA
S2
SIN3 220
250
α-Tubulin
50
KDa
Supplementary Fig 1
