## Supplemental figure 2 for "Isoforms of the transcriptional cofactor SIN3 differentially regulate genes necessary for energy metabolism and cell survival"

### Slide 1
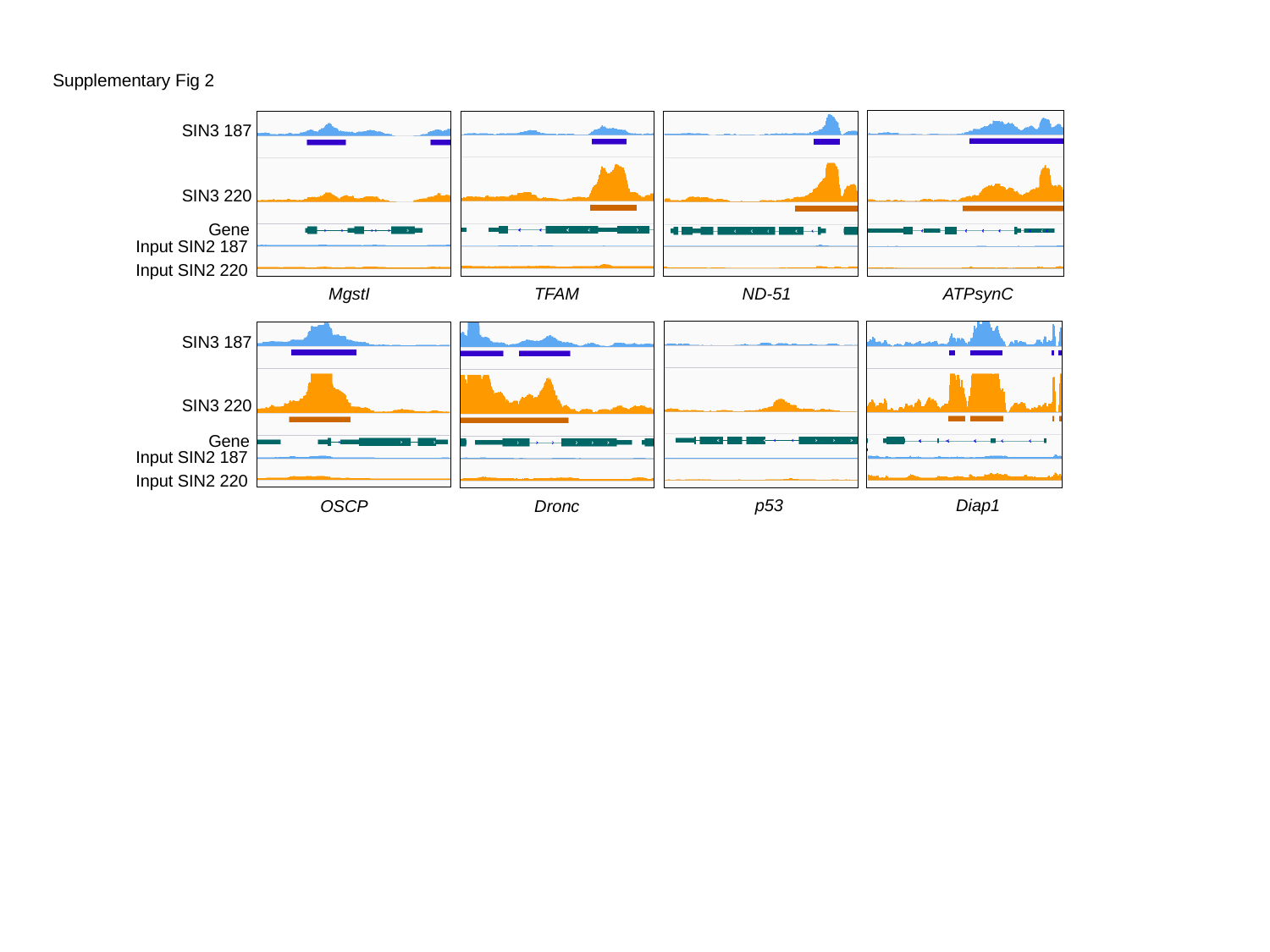

Supplementary Fig 2
SIN3 187
SIN3 220
Gene
Input SIN2 187
Input SIN2 220
MgstI
TFAM
ND-51
ATPsynC
SIN3 187
SIN3 220
Gene
Input SIN2 187
Input SIN2 220
p53
Diap1
Dronc
OSCP
